# Fetal death certificate data quality: A tale of two US counties

**DOI:** 10.1101/136432

**Authors:** Lauren Christiansen-Lindquist, Robert M. Silver, Corette B. Parker, Donald J. Dudley, Matthew A. Koch, Uma M. Reddy, George R. Saade, Robert L. Goldenberg, Carol J. R. Hogue

## Abstract

**Purpose:** Describe the relative frequency and joint effect of missing and misreported fetal death certificate (FDC) data and identify variations by key characteristics.

**Methods:** Stillbirths were prospectively identified during 2006-2008 for a multi-site population-based case-control study. For this study, eligible mothers of stillbirths were not incarcerated residents of DeKalb County, Georgia, or Salt Lake County, Utah, aged > 13 years, with an identifiable FDC. We identified the frequency of missing and misreported (any departure from the study value) FDC data by county, race/ethnicity, gestational age, and whether the stillbirth was antepartum or intrapartum.

**Results:** Data quality varied by item, and was highest in Salt Lake County. Reporting was generally not associated with maternal or delivery characteristics. Reasons for poor data quality varied by item in DeKalb County: some items were frequently missing *and* misreported; however, others were of poor quality due to *either* missing or misreported data.

**Conclusions:** FDC data suffer from missing and inaccurate data, with variations by item and county. Salt Lake County data illustrate that high quality reporting is attainable. The overall quality of reporting must be improved to support consequential epidemiologic analyses for stillbirth, and improvement efforts should be tailored to the needs of each jurisdiction.

**Abbreviations and Acronyms:**

CCC: concordance correlation coefficient
CDC: Centers for Disease Control and Prevention
FDC: Fetal death certificate
NCHS: National Center for Health Statistics
SCRN: Stillbirth Collaborative Research Network

## INTRODUCTION

Although stillbirths (fetal deaths ≤ 20 weeks’ gestation) are now more common than infant deaths,^1^ much less research and attention focuses on reducing stillbirth rates and disparities. Birth and infant death records have been routinely linked since 1980, and have played an important role in maternal and child health epidemiology.^2^ Similar consequential epidemiologic analyses are needed for stillbirth, however the quality of these vital records is lacking.^3–12^

Vital event registration in the United States is decentralized at state or local geographical areas, referred to as jurisdictions,^13^ and vital records data quality may vary by jurisdiction. For example, in 2013, 9.1% of FDCs were missing birth weight (range of jurisdictions: 0.0-42.1%),^14^ compared to only 0.1% of live birth certificates (range of jurisdictions: 0.0–0.9%).^15^16Other FDC variables for which missing data have been a concern include: pregnancy weight gain (70% of records with missing values), gravidity (11%), alcohol and tobacco use during pregnancy (18%), paternal age (74%), and cause(s) of death (69%).^5^ Missing data are also more common for stillbirths compared to neonatal deaths.^11^

Even when data elements are complete, information reported on the FDC is often inaccurate.^3,6,9,12,16–18^ In comparing FDCs to medical records for stillbirths identified through the Wisconsin Stillbirth Service Project,^7^ Greb and colleagues found that FDC-reported sex, birth weight, and gestational age were mostly accurate, but congenital anomalies and cause(s) of death were often misreported. A study conducted in Georgia among stillbirths with implausible birth weight and gestational age values found that approximately one-quarter of these implausible values were due to incorrect reporting.^6^

Previous studies of FDC data quality have focused on data from single jurisdictions. Although there are documented differences in stillbirth rates by race and ethnicity and by gestational age, previous research has not assessed whether data quality varies by these important factors. Further, to our knowledge, the twodimensional aspect of missing and misreported FDC data has not been explored. To address these gaps, we investigated the quality of FDC data by linking records from a population-based case-control study of stillbirth to FDCs. The objective of this study was to assess whether missing and misreported data varied by county of residence, and maternal and/or delivery characteristics, and to describe the joint effect of these biases on FDC data quality.

## MATERIALS AND METHODS

The Stillbirth Collaborative Research Network (SCRN) conducted a multisite, population-based case-control study of mothers of stillbirths and a sample of live births at the time of delivery. Study methods have been previously described.^19^ Enrollment occurred from March 2006 – September 2008 among five clinical sites, each with corresponding catchment areas: Brown University (State of Rhode Island, and Bristol County, MA), Emory University (DeKalb County, GA), University of Texas Medical Branch—Galveston (Galveston and Brazoria Counties, TX), University of Texas Health Science Center—San Antonio (Bexar County, TX), and the University of Utah (Salt Lake County, UT). Hospitals were selected for participation such that at least 90% of all deliveries of catchment area residents would be identified and potentially approached to consent. An effort was made to enroll all eligible residents with stillbirths who were at least 13 years of age, not incarcerated, and identified prior to hospital discharge. Data collection included maternal interview, prenatal care medical chart abstraction, and biological specimens.

For this analysis, records for SCRN-eligible stillbirths identified in Georgia and Utah during the enrollment period were linked to FDCs for all pregnancies with the death of only one fetus. FDCs were not obtainable for participants enrolled in Texas, Massachusetts, and Rhode Island. Study records and FDCs were linked via an iterative deterministic linkage strategy, using varying combinations of portions of the mother’s first and last names, her date of birth, and the date of delivery. For any SCRN stillbirths that did not link, manual searches were conducted using mother’s date of birth, the first and last two letters of her last name, and a review of all FDCs reported within 5 days of the SCRN date of delivery.

The number and proportion of linked records with missing FDC data for maternal and delivery characteristics, prenatal care, and medical risk factors for stillbirth were identified. Due to its detailed data collection protocol, SCRN was used as the gold standard to which FDCs were compared. The source of each SCRN data element is shown in Appendix Table 1. For variables with non-missing data in both FDC and SCRN, we identified the number and proportion of records with misreported FDC data, defined as any departure from the SCRN-recorded value. The accuracy of maternal education was not assessed due to differences in data collection across the data sources.

**Table 1.** Characteristics of 382 residents of DeKalb County, Georgia and Salt Lake County, Utah identified by the Stillbirth Collaborative Research Network study, by county of residence and FDC^a^ linkage status, 2006–2008^b^

| Characteristic<sup>c</sup> | DeKalb County<br>(N = 166) |  |  | Salt Lake County<br>(N = 216) |  |  |
| --- | --- | --- | --- | --- | --- | --- |
|  | FDC Linked | FDC Unlinked | Two-sided exact p-value | FDC Linked | FDC Unlinked | Two-sided exact p-value |
| **Stillbirths** | 126 (75.9) | 40 (24.1) | -- | 208 (96.3) | 8 (3.7) | -- |
| **Method of Linkage** |  |  |  |  |  |  |
| ID 1<sup>d</sup> | 103 (81.7) | -- | -- | 182 (87.5) | -- | -- |
| ID 2<sup>e</sup> | 4 (3.2) | -- | -- | 8 (3.8) | -- | -- |
| ID 3<sup>f</sup> | 5 (4.0) | -- | -- | 8 (3.8) | -- | -- |
| ID 4<sup>g</sup> | 10 (7.9) | -- | -- | 7 (3.4) | -- | -- |
| Manual Search | 4 (3.2) | -- | -- | 3 (1.4) | -- | -- |
| **Maternal Characteristics** |  |  |  |  |  |  |
| Age<sup>h</sup> | 27.0 (6.4) | 26.9 (7.3) | 0.93 | 28.2 (6.4) | 29.1 (6.9) | 0.71 |
| Race/Ethnicity |  |  | 0.18 |  |  | 0.48 |
| Non-Hispanic White | 5 (4.0) | 0 (0) |  | 138 (66.3) | 4 (50.0) |  |
| Non-Hispanic Black | 94 (74.6) | 32 (80.0) |  | 5 (2.4) | 0 (0) |  |
| Hispanic | 17 (13.5) | 2 (5.0) |  | 50 (24.0) | 3 (37.5) |  |
| Other | 10 (7.9) | 6 (15.0) |  | 15 (7.2) | 1 (12.5) |  |
| Education (completed years) |  |  | 0.31 |  |  | 0.61 |
| 0-11 | 23 (18.3) | 5 (12.5) |  | 20 (9.6) | 1 (12.5) |  |
| 12 | 21 (16.7) | 10 (25.0) |  | 41 (19.7) | 2 (25.0) |  |
| 13 or more | 33 (26.2) | 14 (35.0) |  | 91 (43.8) | 2 (25.0) |  |
| Unknown | 49 (38.9) | 11 (27.5) |  | 56 (26.9) | 3 (37.5) |  |
| Mother Married |  |  | 0.08 |  |  | 0.37 |
| Yes | 29 (23.0) | 6 (15.0) |  | 102 (49.0) | 2 (25.0) |  |
| No | 49 (38.9) | 24 (60.0) |  | 51 (24.5) | 3 (37.5) |  |
| Unknown | 48 (38.1) | 10 (25.0) |  | 55 (26.4) | 3 (37.5) |  |
| **Delivery Characteristics** |  |  |  |  |  |  |
| Delivery Year |  |  | < 0.001 |  |  | 0.16 |
| 2006 | 26 (20.6) | 17 (42.5) |  | 52 (25.0) | 0 (0) |  |
| 2007 | 61 (48.4) | 21 (52.5) |  | 90 (43.3) | 6 (75.0) |  |
| 2008 | 39 (31.0) | 2 (0.05) |  | 66 (31.7) | 2 (25.0) |  |
| Gestational Age<sup>h</sup> | 28.1 (6.9) | 26.3 (5.7) | 0.13 | 28.8 (6.9) | 25.1 (6.6) | 0.14 |
| Timing of Death |  |  | 0.14 |  |  | 0.45 |
| Antepartum | 73 (57.9) | 29 (72.5) |  | 138 (66.4) | 4 (50.0) |  |
| Intrapartum | 53 (42.1) | 11 (27.5) |  | 70 (33.7) | 4 (50.0) |  |

<sup>a</sup>Fetal death certificate
<sup>b</sup>Values are n (%) unless otherwise stated
<sup>c</sup>As determined by SCRN
<sup>d</sup>First letter of mother's first name, first two letters of mother's last name, last two letters of mother's last name, mother's date of birth
<sup>e</sup>First letter of mother's first name, first letter of mother's last name, mother's date of birth
<sup>f</sup>First letter of mother's first name, first letter of mother's last name, last letter of mother's last name, date of delivery
<sup>g</sup>Mother's date of birth, date of delivery
<sup>h</sup>Mean (SD)

Fisher’s exact tests were used to evaluate differences in missing and misreported FDC data by SCRN-recorded county of residence, maternal race/ethnicity, and gestational age. Due to differing circumstances surrounding antepartum and intrapartum stillbirths, we also assessed whether data quality was associated with the timing of the death relative to the onset of labor.

Since some discrepancies in the reporting of continuous variables may not be meaningful (e.g. a 2 gram discrepancy in birth weight), we determined whether an individual’s categorization of gestational age and birth weight changed as a result of FDC misreporting, using categories published by the National Center for Health Statistics (NCHS).^1^

Statistical measures of agreement for categorical and continuous variables were calculated using Cohen’s kappa^20^ and Lin’s concordance correlation coefficient (CCC),^21^ respectively. The guidelines of Landis and Koch were used to classify the level of agreement for categorical variables.^22^ The level of agreement for continuous variables was not classified, as the only published guidelines for classifying the CCC were designed for use in a laboratory setting^23^ and are not appropriate for this analysis. We also calculated the sensitivity (the proportion of stillbirths with a given characteristic that were correctly reported on the FDC) as well as the positive predictive value (the proportion of FDCs reporting a particular characteristic that were correctly classified according to SCRN data).

Finally, to describe the joint effect of missing and inaccurate FDC data, we plotted the proportion of FDCs with missing data by the proportion of FDCs with inaccurate data. Data points closest to the origin indicate low levels of both missing and inaccurate data and correspond to variables with the best data quality. Data points further from the origin reflect higher levels of missing and/or misreported data, corresponding to variables with poorer data quality.

This study was reviewed and approved by the Institutional Review Boards of each of the participating sites and the data coordinating center.

## RESULTS

Between March 2006 and September 2008, 166 and 216 pregnancies with a single stillbirth were identified by SCRN in DeKalb and Salt Lake Counties, respectively (Table 1). FDCs were linked to 126 DeKalb stillbirths and 208 Salt Lake stillbirths. Most (n = 285, 85%) FDCs were linked using portions of the mother’s first and last names and her date of birth. There were no statistically significant differences between SCRN stillbirths with and without a linked FDC, except for delivery year among DeKalb County residents.

Missing data were more common for DeKalb than for Salt Lake County stillbirths (Table 2). For DeKalb stillbirths, FDCs frequently lacked maternal education (60%), ethnicity (15%), receipt of prenatal care (37%), number of prenatal care visits (29%), smoking during pregnancy (60%), first pregnancy (19%), and birth weight (16%). Although the DeKalb vital records file included a field to record chronic hypertension, eclampsia, and preeclampsia, these data were missing for all SCRN stillbirths. No variables were missing for more than 10% of Salt Lake County residents. Variables with mostly complete reporting in both counties included maternal race, marital status, county of residence, fetal sex, gestational age, and plurality. After adjusting for county of residence, the frequency of missing data was not associated with maternal race/ethnicity (data not shown). Birth weight and number of prenatal care visits were more likely to be missing for losses occurring at 20–27 weeks’ gestation compared to later losses (birth weight: 14% vs. 5%, p = 0.01; number of prenatal care visits: 21% vs. 12%, p = 0.02). Intrapartum stillbirths were more likely than antepartum stillbirths to be missing information on the receipt of prenatal care (20% vs. 10%, p = 0.01).

**Table 2.** Frequency of missing data for select FDC^a^ data elements for 334 residents of DeKalb County, Georgia and Salt Lake County, Utah identified by the Stillbirth Collaborative Research Network study, by county of residence, 2006–2008^b^

| Characteristic | FDCs with missing value for variable under consideration |  |  |
| --- | --- | --- | --- |
|  | DeKalb County<br>n (%)<br>(N = 126) | Salt Lake County<br>n (%)<br>(N = 208) | Two-sided exact<br>p-value |
| **Maternal Characteristics** |  |  |  |
| Race | 2 (1.6) | 4 (1.9) | 1.0 |
| Ethnicity | 19 (15.1) | 0 (0) | < 0.001 |
| Education (completed years) | 76 (60.3) | 12 (5.8) | < 0.001 |
| Marital status | 0 (0) | 0 (0) | -- |
| County of residence | 0 (0) | 1 (0.5) | -- |
| **Prenatal Care** |  |  |  |
| Received any prenatal care | 47 (37.3) | 0 (0) | < 0.001 |
| Number of prenatal care visits | 37 (29.4) | 18 (8.7) | < 0.001 |
| **Medical Risk Factors for Stillbirth** |  |  |  |
| Smoking during pregnancy | 76 (60.3) | 0 (0) | < 0.001 |
| First Pregnancy<sup>d</sup> | 24 (19.4)<sup>c</sup> | -- | -- |
| Chronic Hypertension | 126 (100) | 0 (0) | < 0.001 |
| Gestational Diabetes<sup>e</sup> | -- | 0 (0) | -- |
| Eclampsia<sup>d</sup> | 126 (100) | -- | -- |
| Preeclampsia<sup>d</sup> | 126 (100) | -- | -- |
| **Delivery Characteristics** |  |  |  |
| Sex | 3 (2.4) | 2 (1.0) | 0.37 |
| Gestational Age | 0 (0) | 0 (0) | -- |
| Birth Weight | 20 (15.9) | 13 (6.3) | 0.007 |
| Plurality | 4 (3.2) | 0 (0) | 0.02 |

<sup>a</sup>Fetal death certificate
<sup>b</sup>Values are n(%)
<sup>c</sup>Two women had FDC values indicating “not applicable” for this variable and have been removed from the denominator
<sup>d</sup>Variable not collected on Salt Lake County FDCs
<sup>e</sup>Variable not collected on DeKalb County FDCs

When data were present, they were more likely to be misreported for DeKalb than for Salt Lake County stillbirths (Table 3). Variables most frequently misreported were: number of prenatal care visits (DeKalb: 78%, Salt Lake: 52%), gestational age (DeKalb: 54%, Salt Lake: 21%), birth weight (DeKalb: 49%, Salt Lake: 13%), and maternal race (DeKalb: 19%, Salt Lake: 11%). Variables with moderate levels of misreporting were marital status, county of residence (DeKalb), and smoking during pregnancy (DeKalb). Variables most accurately reported were maternal ethnicity, receipt of prenatal care, chronic hypertension (Salt Lake), gestational diabetes (Salt Lake), fetal sex, and plurality. Misreporting of FDC data elements was not associated with maternal race/ethnicity, gestational age, or the timing of the death (data not shown).

**Table 3.** Frequency of misreported FDC^a^ information for select data elements for residents of DeKalb County, Georgia and Salt Lake County, Utah identified by the Stillbirth Collaborative Research Network study, by county of residence, 2006–2008^b^

| Characteristic | FDCs with misreported value for variable under consideration |  |  |
| --- | --- | --- | --- |
|  | DeKalb County<br>n (%) | Salt Lake County<br>n (%) | Two-sided exact<br>p-value |
| **Maternal Characteristics** |  |  |  |
| Race (n = 328) | 23 (18.6) | 22 (10.8) | 0.07 |
| Ethnicity (n = 315) | 2 (1.9) | 7 (3.4) | 0.72 |
| Marital Status (n = 231) | 6 (7.7) | 7 (4.6) | 0.37 |
| County of Residence (n = 333) | 11 (8.7) | 3 (1.5) | 0.003 |
| **Prenatal Care** |  |  |  |
| Received any Prenatal Care (n = 201) | 1 (2.1) | 3 (2.0) | 1.0 |
| Number of Prenatal Care Visits (n = 192) | 39 (78.0) | 74 (52.1) | 0.001 |
| **Medical Risk Factors for Stillbirth** |  |  |  |
| Smoking During Pregnancy (n = 176) | 4 (15.4) | 7 (4.7) | 0.06 |
| First Pregnancy (n = 65)<sup>c</sup> | 4 (6.2) | -- | -- |
| Chronic Hypertension (n = 147)<sup>d</sup> | -- | 2 (1.4) | -- |
| Gestational Diabetes (n = 160)<sup>e</sup> | -- | 6 (3.8) | -- |
| **Delivery Characteristics** |  |  |  |
| Sex (n = 238) | 1 (1.3) | 1 (0.6) | 1.0 |
| Gestational Age (n = 334) | 68 (54.0) | 43 (20.7) | < 0.001 |
| Birth Weight (n = 211) | 32 (48.5) | 19 (13.1) | < 0.001 |
| Plurality (n = 330) | 0 (0) | 0 (0) | -- |

<sup>a</sup>Fetal death certificate
<sup>b</sup>Since the SCRNs and vital records data availability differ for each variable, the total number of women with non-missing data in both sources is shown in parentheses
<sup>c</sup>Variable not collected in Salt Lake County
<sup>d</sup>Variable completely missing in DeKalb County
<sup>e</sup>Variable not collected in DeKalb County

Misreported gestational age and birth weight resulted in changes in NCHS categorizations for many stillbirths (Table 4). As a result of misreporting on the FDC, 64 (58%) of the 111 stillbirths with a misreported FDC value for week of gestational age changed NCHS gestational age categories, and 9 (18%) of 51 stillbirths with an inaccurate FDC value for birth weight changed NCHS birth weight categories. No differences in NCHS categorization of gestational age or birthweight were observed by county of residence (data not shown).

**Table 4.** Changes in NCHS gestational age or birth weight group membership among residents of DeKalb County, Georgia and Salt Lake County, Utah identified by the Stillbirth Collaborative Research Network study with misreported information on the FDC^a^, 2006–2008

|  |  | **FDC Reporting of Gestational Age** |  |  |
| --- | --- | --- | --- | --- |
| **SCRN Gestational Age (weeks)** | **n (%)** | **Classified at Earlier Gestations**<br>**n (%)** | **No Change in Group Membership**<br>**n (%)** | **Classified at Later Gestations**<br>**n (%)** |
| All Gestational Ages | 111 (100) | 27 (24.3) | 47 (42.3) | 37 (33.3) |
| NCHS Gestational Age Groups |  |  |  |  |
| 20-23 | 37 (33.3) | 3 (8.1) | 20 (54.1) | 14 (37.8) |
| 24-27 | 19 (17.1) | 4 (21.1) | 10 (52.6) | 5 (26.3) |
| 28-31 | 14 (12.6) | 9 (64.3) | 3 (21.4) | 2 (14.3) |
| 32-33 | 5 (4.5) | 2 (40.0) | 2 (40.0) | 1 (20.0) |
| 34-36 | 20 (18.0) | 4 (20.0) | 8 (40.0) | 8 (40.0) |
| 37-39 | 11 (9.9) | 2 (18.2) | 4 (36.4) | 5 (45.5) |
| 40 | 2 (1.8) | 1 (50.0) | 0 (0) | 1 (50.0) |
| 41 | 3 (2.7) | 2 (66.7) | 0 (0) | 1 (33.3) |
| ≥ 42<sup>b</sup> | 0 (0) | NA | NA | NA |
| 20-27 | 56 (50.5) | 3 (5.4) | 43 (76.8) | 10 (17.9) |
| ≥ 28 | 55 (49.6) | 11 (20.0) | 44 (80.0) | NA |
|  |  | **FDC Reporting of Birth Weight** |  |  |
| **SCRN Birth Weight (grams)** | **n (%)** | **Classified at Lower Birth Weights**<br>**n (%)** | **No Change in Group Membership**<br>**n (%)** | **Classified at Greater Birth Weights**<br>**n (%)** |
| All Birth Weights | 51 (100) | 3 (5.9) | 42 (82.4) | 6 (11.8) |
| NCHS Birth Weight Groups |  |  |  |  |
| < 500 | 15 (29.4) | NA | 12 (80.0) | 3 (20.0) |
| 500 – 749 | 7 (13.7) | 0 (0) | 5 (71.4) | 2 (28.6) |
| 750 – 999 | 6 (11.8) | 0 (0) | 6 (100) | 0 (0) |
| 1,000 – 1,249 | 3 (5.9) | 1 (33.3) | 2 (66.7) | 0 (0) |
| 1,250 – 1,499 | 2 (3.9) | 0 (0) | 2 (100) | 0 (0) |
| 1,500 – 1,999 | 5 (9.8) | 1 (20.0) | 4 (80.0) | 0 (0) |
| 2,000 – 2,499 | 6 (11.8) | 0 (0) | 6 (100) | 0 (0) |
| 2,500 – 2,999 | 4 (7.8) | 1 (25.0) | 2 (50.0) | 1 (25.0) |
| 3,000 – 3,499 | 2 (3.9) | 0 (0) | 2 (100) | 0 (0) |
| 3,500 – 3,999 | 1 (2.0) | 0 (0) | 1 (100) | 0 (0) |
| ≥ 4,000<sup>c</sup> | 0 (0) | NA | NA | NA |

<sup>a</sup>Fetal death certificate
<sup>b</sup>No SCRN cases with gestational age ≥ 42 weeks with an inaccurate FDC report of gestational age
<sup>c</sup>No SCRN cases with a birth weight ≥ 4,000 grams with an inaccurate FDC report of birth weight

The majority of FDCs reported a gestational age within one week of the SCRN value (DeKalb: 71.4%, Salt Lake: 95.2%). Some FDCs (DeKalb: 15.9%, Salt Lake: 2.4%) reported a gestational age that differed from the SCRN value by 4 weeks or more. Similarly, a majority of FDCs reported a birth weight within 9 grams of the SCRN value (DeKalb: 68.2%, Salt Lake: 90.3%); however a number of FDCs reported a birth weight that was different from the SCRN value by 51 grams or more (DeKalb: 15.2%, Salt Lake: 3.4%). The majority of FDCs reported the number of prenatal care visits within one visit of the SCRN value (DeKalb: 54.0%, Salt Lake: 74.6%).

Statistical measures of agreement between SCRN and vital records are shown in Appendix Tables 2a and 2b. Agreement was almost perfect for maternal ethnicity, marital status, fetal sex, and plurality in both counties. In DeKalb agreement was only moderate for race, and smoking during pregnancy. The sensitivities and positive predictive values largely correspond to the levels of agreement noted above. Although the positive predictive value for maternal smoking was 100% in DeKalb County, only one-third of SCRN-identified smokers had this designation on the fetal death certificate.

The joint effect of missing and inaccurate data is shown in Figures la-lc. The proportion of records with inaccurate values was calculated among records with non-missing data. In DeKalb County, variables with the best overall data quality were fetal sex, plurality, and marital status. Primarily because of missing values, data quality was worse for ethnicity, gravidity, receipt of prenatal care, smoking, eclampsia, and preeclampsia.

Gestational age and maternal race were generally not missing but were often inaccurate, whereas birth weight and number of prenatal care visits were more often missing and, when given, inaccurate. In Salt Lake County, variables with the best overall data quality included plurality, fetal sex, receipt of prenatal care, maternal ethnicity, marital status, county of residence, smoking, and gestational diabetes. Data quality was worse for birth weight (due to missing data); gestational age and maternal race (due to inaccurate data); and number of prenatal care visits (due to both missing and inaccurate data).

## DISCUSSION

This study reflects FDC data quality for stillbirths occurring to women who were eligible for enrollment by SCRN and were issued an FDC from March 2006 – September 2008. Substantial differences in data quality by maternal and delivery characteristics were not observed; however, data quality was highly associated with mother’s county of residence. In both counties, fetal sex, plurality, and marital status tended to be reported both completely and accurately. Additionally, Salt Lake County had high quality data for receipt of prenatal care, maternal ethnicity, county of residence, smoking, and gestational diabetes. In both counties, the quality of reporting of maternal race and gestational age suffered more from inaccurate (rather than missing) values. The quality of several variables in DeKalb County was affected more by missing values.

Both counties had near complete reporting of gestational age, but these values were frequently misreported, with misclassification of stillbirths in the NCHS gestational age categories for 58% of stillbirths with incorrect FDC values. Delivery facilities receive guidance for reporting gestational age in two forms: the NCHS Guide to Completing the Facility Worksheets for the Certificate of Live Birth and Report of Fetal Death^24^ (Guide) and the Facility Worksheet for the Report of Fetal Death^25^ (Worksheet). Since there may be a period of time during which a fetus is still in utero, but no longer alive, gestational age reporting is more complicated for stillbirths than for live births. Despite this difference, the Guide does not differentiate reporting of the obstetric estimate of gestation for live births and stillbirths. Additionally, these resources provide conflicting information for reporting of gestational age. The Guide suggests that the delivery attendant *could* use the mother’s date of last menstrual period and the date of delivery, whereas the Worksheet indicates that this information *should not* be used to provide an estimate of the gestation at delivery. A revision of these instructions may help to improve data quality, and instructions for reporting the obstetric estimate of gestation should distinguish between stillbirths and live births.

NCHS recommends that data be collected using the most reliable source; some variables are derived from medical records, while others come from self-report. Women are asked to provide information regarding their race, ethnicity, education, marital status, and smoking during pregnancy on the Patient’s Worksheet for the Report of Fetal Death.^26^ With the exception of marital status, all of these variables were frequently missing or misreported in DeKalb County; specifically, both maternal education and smoking were missing for 76 women, suggesting that they did not complete the Patient’s Worksheet. While maternal race was rarely missing on the FDC, it was incorrect for 19% of DeKalb residents. Maternal ethnicity was missing for 15% of women in DeKalb County, but was only misreported for 2% of those with a non-missing value. If the Patient’s Worksheet was not available, maternal race and ethnicity may have been recorded by hospital staff, with potential for error. It is unknown whether the patients chose not to complete the form, or if they were not asked to do so. Although hospital staff may be concerned about burdening patients with this administrative task after a stillbirth, women should still be given the opportunity to answer these demographic questions. Only 17% of women approached to participate in the SCRN study refused.^19^ While there were benefits to the individual for participating (i.e. autopsy results and referral for grief support), women who have experienced a stillbirth have expressed willingness to answer questions about their experiences when the only benefit was that researchers might learn more about stillbirth and prevent future losses.^27^ For these reasons, it is likely that women would be willing to answer these demographic questions if they were approached in an appropriate way. Hospital staff should receive training on the importance of collecting these data as well as techniques for approaching bereaved mothers to request this information in a sensitive manner.

### Limitations

Although SCRN data were considered the gold standard, there may have been instances where the FDC had reported the correct value. We believe this to be a rare occurrence as SCRN collected information for all study enrollees via medical record abstraction and maternal interview prior to hospital discharge. Additionally, this study included all women who were eligible for the case-control study; however not all women consented to participate in all portions of the study. For this reason, we were missing data items for comparison to the vital record for some women. Also, stillbirths not identified by SCRN were not included in this analysis. To the extent that SCRN-missed stillbirths differ from those in our sample, our findings may not represent the data quality for these catchment areas. Finally, these data do not provide insight into the reasons for the differences in data quality between the two counties. Efforts are underway to investigate fetal death reporting in Georgia to better understand the causes of poor reporting.

This study contributes to the body of literature regarding FDC data quality by examining: data from a population-based sample in two disparate counties; whether FDC data quality is associated with maternal and delivery characteristics; and the joint effect of missing and misreported data. The high levels of missing and misreported data observed in DeKalb County confirm previous studies. Additionally, our study demonstrates that data quality is not associated with maternal or delivery characteristics, but rather with county of residence, which corresponds to the vital statistics reporting jurisdiction (i.e. State). Within a given jurisdiction, certain groups do not appear to have worse data quality than others; however, to the extent that the distribution of these factors varies across jurisdictions, national stillbirth rates stratified by these factors may be impacted.

A recent study showed that facility-reported barriers to fetal death reporting were associated with completeness and accuracy of FDC data in New York City.^10^ This, along with our results, suggests that barriers to reporting likely vary by jurisdiction, and interventions tailored to the needs of each jurisdiction might be necessary.

Our findings from Salt Lake County indicate that high quality FDC reporting is possible. Financial and technical assistance resources are needed to facilitate the collection of timely and accurate fetal death data.

Efforts aimed at improving FDC data could include: clarifying instructions for reporting gestational age; training of health providers and hospital staff on the importance of collecting high quality data (including the importance of the Patient Worksheet); performing regular audits of data accuracy; linkage with electronic medical records; and revisiting and revising the cause of death, as needed, after all testing, including autopsy, is complete.

## Acknowledgments and Funding

This research was supported by grant funding from the Eunice Kennedy Shriver National Institute of Child Health and Human Development (NICHD): U10-HD045925 Emory University; U10-HD045944 University of Utah Health Sciences Center; and U01-HD045954 RTI International, RTP; and NIH training grant T32HD052460 Emory University. This project was also supported by the Health Resources and Services Administration (HRSA) of the U.S. Department of Health and Human Services (HHS) under the Maternal and Child Health Epidemiology Doctoral Training Program (Grant #T03MC07651). This content and conclusions are those of the author and should not be construed as the official position or policy of, nor should any endorsements be inferred by HRSA, HHS or the U.S. Government.

**Appendix Table 1.** Source of gold standard data used for comparison to fetal death certificates

| Data Element | Source of Gold Standard Data |  |
| --- | --- | --- |
|  | Medical Records | Maternal Interview<sup>a</sup> |
| **Maternal Characteristics** |  |  |
| Race |  | X<sup>b</sup> |
| Ethnicity |  | X<sup>b</sup> |
| Marital Status |  | X |
| County of Residence | X |  |
| **Prenatal Care** |  |  |
| Received any Prenatal Care |  | X |
| Number of Prenatal Care Visits | X |  |
| **Medical Risk Factors for Stillbirth** |  |  |
| Smoking During Pregnancy |  | X<sup>c</sup> |
| First Pregnancy | X<sup>d</sup> |  |
| Chronic Hypertension | X |  |
| Gestational Diabetes | X |  |
| **Delivery Characteristics** |  |  |
| Sex | X |  |
| Gestational Age | X<sup>e</sup> |  |
| Birth Weight | X |  |
| Plurality | X |  |

<sup>a</sup>Collected in the hospital shortly after delivery
<sup>b</sup>Where available, otherwise taken from medical records
<sup>c</sup>Maternal interview was selected as the gold standard source for smoking during pregnancy in accordance with the NCHS instructions for completing the fetal death certificate
<sup>d</sup>Where available, otherwise taken from maternal interview
<sup>e</sup>Determined using documentation for assisted reproduction or via a combination of ultrasound measurements and the first date of the last menstrual period

**Appendix Table 2a.** Statistical measures of agreement between select Fetal Death Certificate data elements and data collected by the Stillbirth Collaborative Research Network for residents of DeKalb County, Georgia, 2006–2008

| Characteristic | Measure of Agreement<sup>a</sup><br>(95% Confidence Interval) | Level of Agreement<sup>b</sup> | Sensitivity<sup>c</sup> | Positive Predictive Value<sup>c</sup> |
| --- | --- | --- | --- | --- |
| **Maternal Characteristics** |  |  |  |  |
| Race<sup>d</sup> | 0.51 (0.39, 0.62) | Moderate | 98.9%<sup>g</sup> | 93.0%<sup>h</sup> |
| Ethnicity<sup>d</sup> | 0.93 (0.83, 1.00) | Almost perfect | 93.8% | 93.8% |
| Marital Status<sup>d</sup> | 0.84 (0.71, 0.96) | Almost perfect | i | i |
| **Prenatal Care (PNC)** |  |  |  |  |
| Did not receive PNC<sup>d</sup> | 0.79 (0.39, 1.00) | Substantial | 100% | 66.7% |
| Number of PNC Visits<sup>e</sup> | 0.54 (0.36, 0.67) | -- | -- | -- |
| **Medical Risk Factors for Stillbirth** |  |  |  |  |
| Smoked During Pregnancy<sup>d</sup> | 0.43 (0.01, 0.86) | Moderate | 33.3% | 100% |
| First Pregnancy<sup>d</sup> | 0.83 (0.66, 0.99) | Almost perfect | 86.7% | 86.7% |
| Chronic Hypertension | f | -- | f | f |
| Gestational Diabetes | f | -- | f | f |
| **Delivery Characteristics** |  |  |  |  |
| Sex<sup>d</sup> | 0.97 (0.92, 1.00) | Almost perfect | 100% | 97.8% |
| Gestational Age<sup>e</sup> | 0.89 (0.85, 0.92) | -- | -- | -- |
| Birth Weight<sup>e</sup> | 0.90 (0.84, 0.94) | -- | -- | -- |
| Plurality<sup>d</sup> | 1.00 (1.00, 1.00) | Almost perfect | 100% | 100% |

<sup>a</sup>Kappa reported for categorical variables and concordance correlation reported for continuous variables
<sup>b</sup>Using Landis and Koch classification for categorical variables
<sup>c</sup>Not reported for continuous variables
<sup>d</sup>Kappa
<sup>e</sup>Concordance correlation coefficient
<sup>f</sup>Variable not collected in DeKalb County
<sup>g</sup>P(reported as Black on vital records | mother reporting Black race to SCRn)
<sup>h</sup>P(mother reporting Black race to SCRn | reported as Black on vital records)
<sup>i</sup>Did not calculate sensitivity and positive predictive value due to missing SCRn data

**Appendix Table 2b.** Statistical measures of agreement between select Fetal Death Certificate data elements and data collected by the Stillbirth

| Characteristic | Measure of Agreement<sup>a</sup><br>(95% Confidence Interval) | Level of Agreement<sup>b</sup> | Sensitivity<sup>c</sup> | Positive Predictive Value<sup>c</sup> |
| --- | --- | --- | --- | --- |
| **Maternal Characteristics** |  |  |  |  |
| Race<sup>d</sup> | 0.74 (0.64, 0.84) | Substantial | 75.0%<sup>g</sup> | 100%<sup>h</sup> |
| Ethnicity<sup>d</sup> | 0.91 (0.84, 0.97) | Almost perfect | 90.0% | 95.7% |
| Marital Status<sup>d</sup> | 0.90 (0.82, 0.97) | Almost perfect | i | i |
| **Prenatal Care (PNC)** |  |  |  |  |
| Did not receive PNC<sup>d</sup> | 0.66 (0.29, 1.00) | Substantial | 75.0% | 60% |
| Number of PNC Visits<sup>e</sup> | 0.64 (0.55, 0.72) | -- | -- | -- |
| **Medical Risk Factors for Stillbirth** |  |  |  |  |
| Smoked During Pregnancy<sup>d</sup> | 0.80 (0.66, 0.94) | Substantial | 81.0% | 85% |
| First Pregnancy | f | -- | -- | -- |
| Chronic Hypertension<sup>d</sup> | 0.87 (0.69, 1.00) | Almost perfect | 77.8% | 100% |
| Gestational Diabetes<sup>d</sup> | 0.61 (0.32, 0.89) | Substantial | 55.6% | 71.4% |
| **Delivery Characteristics** |  |  |  |  |
| Sex<sup>d</sup> | 0.99 (0.96, 1.00) | Almost perfect | 100% | 98.8% |
| Gestational Age<sup>e</sup> | 0.98 (0.97, 0.98) | -- | -- | -- |
| Birth Weight<sup>e</sup> | 0.98 (0.97, 0.99) | -- | -- | -- |
| Plurality<sup>d</sup> | 1.00 (1.00, 1.00) | Almost perfect | 100% | 100% |

Collaborative Research Network for residents of Salt Lake County, Utah, 2006-2008
<sup>a</sup>Kappa reported for categorical variables and concordance correlation reported for continuous variables
<sup>b</sup>Using Landis and Koch classification for categorical variables
<sup>c</sup>Not reported for continuous variables
<sup>d</sup>Kappa
<sup>e</sup>Concordance correlation coefficient
<sup>f</sup>Variable not collected in Salt Lake County
<sup>g</sup>P(reported as Black on vital records | mother reporting Black race to SCRNM)
<sup>h</sup>P(mother reporting Black race to SCRNM | reported as Black on vital records)

